# An open source plant kinase chemogenomics set

**DOI:** 10.1101/2022.06.18.496431

**Authors:** Maria Florencia Ercoli, Priscila Zonzini Ramos, Rashmi Jain, Joseph Pilotte, Oliver Xiaoou Dong, Ty Thompson, Carrow I. Wells, Jonathan M. Elkins, Aled M Edwards, Rafael M. Couñago, David H. Drewry, Pamela C. Ronald

**Affiliations:** Centro de Química Medicinal (CQMED), Centro de Biologia Molecular e Engenharia Genética (CBMEG), Universidade Estadual de Campinas (UNICAMP), Campinas, SP, 13083-875, Brazil; Centre for Medicines Discovery, University of Oxford, Old Road Campus Research Building, Roosevelt Drive, Oxford, OX3 7DQ, UK; Department of Plant Pathology and the Genome Center, University of California, Davis, CA, USA; Structural Genomics Consortium (SGC), UNC Eshelman School of Pharmacy, University of North Carolina at Chapel Hill (UNC-CH), Chapel Hill, NC 27599, USA; Division of Chemical Biology and Medicinal Chemistry, UNC Eshelman School of Pharmacy, UNC-CH, Chapel Hill, NC 27599, USA; Structural Genomics Consortium, University of Toronto, Toronto, Canada

**Keywords:** *Oryza sativa*, plant kinases, ligation-independent cloning, protein production, compound screening, thermal shift assay

## Abstract

129 protein kinases, selected to represent the diversity of the rice (*Oryza sativa*) kinome, were cloned and tested for expression in *E. coli*. 40 of these rice kinases were purified and screened using differential scanning fluorimetry (DSF) against 627 diverse kinase inhibitors, with a range of structures and activities targeting diverse human kinases. 37 active compounds were then tested for their ability to modify primary root development in Arabidopsis. Of these, 14 compounds caused a significant reduction of primary root length and two slightly increased root elongation compared with control plants. Two inhibitory compounds bind to the predicted orthologue of Arabidopsis PSKR1, one of two receptors for PSK, a small sulfated peptide that positively controls root development. Inhibition could not be rescued by the exogenous addition of the PSK peptide, suggesting that chemical treatment may inhibit both PSKR1 and its closely related receptor PSKR2. Of the compounds acting as root growth inhibitors in Arabidopsis, six conferred the same effect in rice. Compound RAF265 (CHIR-265), previously shown to bind the human kinase BRAF (B-Raf proto-oncogene, serine/threonine kinase), also binds to nine highly conserved rice kinases tested. The binding of human and rice kinases to the same compound suggests that human kinase inhibitor sets will be useful for dissecting the function of plant kinases.

## Introduction

Protein phosphorylation is the most common form of posttranslational modification used in signal transduction by eukaryotic cells. In plants, protein kinases regulate key biological responses, such as hormone levels, metabolism, morphology, growth, and development (Deprost et al., 2007; Wang et al., 2010; Garcia et al., 2012; Marshall et al., 2012; Bhargava and Sawant, 2013; Osakabe et al., 2013; Danquah et al., 2014; Wierzba and Tax, 2013; Wu and Cheng, 2014; Todaka et al., 2015). As in other eukaryotes, protein kinases constitute one of the largest protein families within plant genomes. In rice (*Oryza sativa*), there are about 1,500 genes that encode for recognizable protein kinase domains (∼3.5% of the rice genome), the vast majority of which remain uncharacterized (Manning et al., 2002; Goff et al., 2002; Yu et al., 2002; Yamamoto et al., 2012; Chandran et al., 2016).

Genetic approaches, such as gene knockouts, have successfully identified plant kinases that mediate important traits but can be confounded by the fact that many plant genes have functionally redundant paralogues (Hicks and Raikhel, 2009). As a result, more than 40% of the genes in a plant genome are “invisible” to single knockout genetic screens. In addition, genes that cause lethality when knocked out cannot be discovered in these screens. This gap presents an opportunity for basic and applied science.

An alternative approach to genetic manipulation is to use a chemical biology strategy based on small molecule modulators of protein kinase function (Hicks and Raikhel, 2009, 2014). Protein kinases share similar ATP-binding sites, and it is not uncommon for small molecule kinase inhibitors to be active against multiple, closely related kinases, suggesting that a kinase inhibitor may chemically “knockout” paralogues or even small families of kinases. Thus, using sets of carefully selected, well-characterized kinase inhibitors that cover most of an organism’s kinome in phenotypic assays allows the observed biological effect to be narrowed down to a small number of kinases (Uitdehaag et al., 2012). For human proteins, the construction of such a kinase chemogenomic set has allowed this strategy to successfully illuminate new biology and discover new therapeutic opportunities (Al-Ali et al., 2015; Jones and Bunnage, 2017; Burdova et al., 2019; Wells et al., 2021). A similar approach has also been used to perform cost-effective, chemistry-based synthetic lethal screens in plants (Hicks and Raikhel, 2009; Xuan et al., 2013; Hicks and Raikhel, 2014). Nevertheless, the lack of well-characterized small molecule reagents has limited the exploration of plant kinomes.

Establishing a well-characterized, broadly distributed Rice Kinase Chemogenomic Set would allow the scientific community to explore the function of rice kinases and deepen our understanding of plant signaling pathways. This endeavor would require the recombinant production of soluble, active rice kinases, the establishment of high-throughput screening (HTS) assays to identify small molecule ligands from libraries of compounds, and iterative chemistry to optimize compound selectivity profiles. These compounds would then be used in phenotypic screens to investigate the biological impact of modulating the function(s) of the target kinase(s). The on-target activity of inhibitors that confer interesting phenotypes could then be verified via chemoproteomics and further validated using genetic tools, such as the creation of rice knockout lines (Huber and Superti-Furga, 2016). Broad distribution would allow the community to use this compound set in a range of phenotypic assays relevant to different facets of plant biology.

Importantly, the conservation of the overall protein kinase architecture, biochemical activity, and ATP-binding site across distantly-related species should allow the knowledge, protocols, assays, and reagents obtained during the development of the human kinase chemogenomic set to be used in the establishment of a similar set of reagents for rice kinases. Indeed, it is now well-established that small molecule inhibitors originally designed for human kinases are also active against kinases from unrelated organisms, such as eukaryotic parasites and plants (Peña et al., 2015; Aquino et al., 2017; Alam et al., 2019). Likewise, the strategy to combine available structural information with high-throughput cloning adopted by structural genomics initiatives to expedite the recombinant production of soluble, active human proteins (Savitsky et al., 2010) has also been shown effective for plant proteins (Tosarini et al., 2018). Finally, HTS assays used to identify ligands for human proteins (Niesen et al., 2007) have been applied with success for plant protein kinases (Aquino et al., 2017).

Here we established the groundwork for the creation of a Rice Kinase Chemogenomic Set and identified a previously unknown connection between 16 compounds and primary root length. We also show that one compound, previously shown to bind the human kinase BRAF (B-Raf proto-oncogene, serine/threonine kinase), also binds at least to nine rice kinases. Our data thus suggests that the methods used for the generation of the human kinase chemogenomics set are readily applicable to dissecting kinase function in plants. Further, we show that small molecule kinase inhibitors can be used to identify new biological processes, contributing to the development of knowledge that will be of interest to the wider plant science community.

## Results

### Selection of protein kinases

The rice genome has 1,467 genes encoding a recognizable protein kinase domain. These can be divided into 63 distinct kinase families belonging to six kinase groups (AGC, CAMK, CK1, CMGC, STE and TKL) based on sequence identity levels as established by the rice kinase phylogenomics database (Dardick et al., 2007; Jung et al., 2015) (Figure 1). To select a representative set of protein kinase genes from the rice genome, we first checked expression values of these genes in 21 available RNA-Seq libraries from the Rice Genome Annotation Project database containing data from samples collected from various rice tissues during different developmental stages or under various biotic and abiotic stresses (Kawahara et al., 2013). We selected 975 genes having expression levels <u>>=2.0 from this analysis.</u>

**Figure 1.**
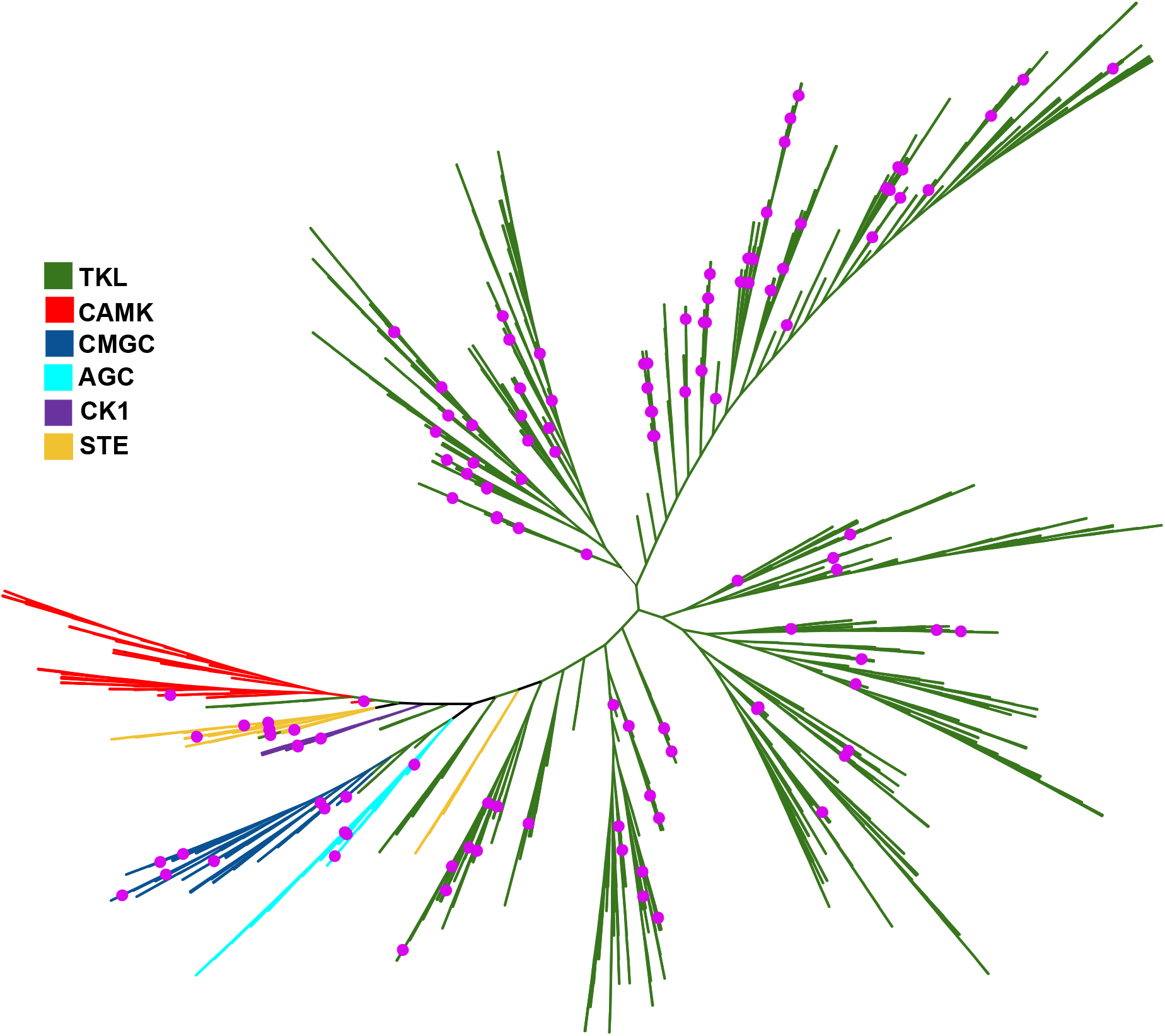
A phylogenetic tree showing the rice kinases selected for this study (Pink dots). The rice kinome contains 1,467 proteins that are classified into six kinase groups (TKL (Tyrosine Kinase-Like), Green; CAMK (Ca2+/calmodulin-dependent protein kinase), Red; CMGC (cyclin-dependent kinase (<u>C</u>DK), mitogen-activated protein kinase (<u>M</u>APK), glycogen synthase kinase (<u>G</u>SK) and CDC-like kinase (<u>C</u>LK)), Blue; AGC (<u>A</u>MP dependent kinases (PKA), c<u>G</u>MP-dependent kinases, and the diacylglycerol-activated/phospholipid-dependent kinase PK<u>C)</u>, Cyan; CK1 (Casein kinase 1), Purple; STE (<u>Sterile serine/threonine kinases)</u>, Saffron). The phylogenetic tree was constructed using the unweighted neighbor-joining method and drawn using Interactive Tree Of Life (iTOL) v5 online tool (Letunic & Bork, 2021).

We next employed RICENet v2, a probabilistic gene network to enrich for trait-associated genes amongst the selected 975 rice protein kinase-encoding genes (Lee et al., 2011, 2015). This analysis resulted in the selection of 141 kinase-encoding genes representing 45 out of the 63 kinase families predicted to participate in independent pathways. Then, we selected one kinase-encoding gene from each of the remaining 18 kinase families to ensure that at least each kinase family was represented by at least one member. Finally, we also included in our set three well-studied kinase-encoding genes: the kinase domain of the rice disease resistance gene XA21(AAC49123) (Song et al., 1997), the XA21-coreceptor (OsSERK2, LOC_Os04g38480) (Chen et al., 2014), and a histidine kinase (LOC_Os06g44410) (Taylor et al., 2021) known to regulate rice root development. Thus, the initially selected set consisted of 162 genes. We further predicted domain information of these kinases using Pfam (Mistry et al., 2021). Out of the 162 selected rice genes, we removed 15 whose gene products lacked a predicted full kinase domain and thus are unlikely to bind inhibitors. Among the remaining genes, we could not obtain synthetic DNA for 18 due to gene synthesis failure (including the histidine kinase LOC_Os06g44410). Following subtraction of these genes, the final set consisted of 129 rice kinases representing diversity within the rice kinome that were successfully synthesized (Figure 1, Supplemental Table S1).

### Recombinant production of selected rice protein kinases

Heterologous expression of eukaryotic genes in a bacterial host may lead to the production of insoluble or inactive recombinant protein. Here we adopted a high-throughput, protein structure-based strategy to quickly identify protein constructs that can be recombinantly produced in a soluble form in *Escherichia coli* (Savitsky et al., 2010; Tosarini et al., 2018). For each of the 129 selected rice protein kinase genes, we designed an average of four different constructs for expression of the isolated kinase domain with varying N- and C-termini. Construct design was based on the best matches from the Protein Data Bank (PDB) for each of the selected rice kinases, identified using the PSIPRED server (Buchan and Jones, 2019). DNA fragments representing each of these kinase domain truncations were obtained via PCR using the appropriate set of synthetic DNA template and oligonucleotide primers (see Supplemental Data Set 1). Amplicons were cloned via ligation-independent cloning into a pET28-based expression vector which added a cleavable 6xHis tag to the N-terminus of the recombinant protein (Aslanidis and de Jong, 1990; Stols et al., 2002; Strain-Damerell et al., 2014). In total, 515 constructs, representing all 129 selected rice kinase-encoding genes, were successfully cloned (see Supplemental Data Set 1).

Soluble recombinant production of all 515 rice kinase constructs in two different *E. coli* strains was evaluated using small-scale test expression (1 mL cultures) followed by purification via ion metal affinity chromatography (IMAC, facilitated by the presence of the N-terminal 6xHis tag in the recombinant protein) from clarified cell lysates. IMAC eluates were visualized by denaturing polyacrylamide gel electrophoresis (SDS-PAGE). These analyses revealed that 286 of the 515 constructs (55.5%) could be purified from clarified cell lysates - as indicated by the presence of a protein band of the expected molecular weight (see Supplemental Data Set 1; Supplemental Figure S1). Overall, we could detect the soluble production of 85 out of the 129 selected rice protein kinases (66%). 40 of these protein kinases were then purified in milligram scale for chemical screening studies.

### Ligand identification

To identify ligands for the purified rice kinases from a library of commercially-available human kinase inhibitors, we used a thermal-stability assay (Differential Scanning Fluorimetry, DSF). This assay is based on the ability of a ligand to stabilize a target protein and increase its temperature-induced unfolding midpoint (*T*_m_) compared to a no-ligand control (reported as a Δ*T*_m_). DSF has been extensively employed to assess binding of compounds to target protein kinases and to estimate compound promiscuity (Fedorov et al., 2012; Elkins et al., 2016). Compound library selection took into account three main criteria. First, all compounds used here are readily-available from commercial vendors. This makes it easy to obtain compounds for follow-up phenotypic assays in plants, which are likely to use large quantities of material. Secondly, the 627 compounds included in our library have a wide range of chemical scaffolds. As the development of plant kinase inhibitors is still in its infancy, we opted to use a library with a large chemical diversity. Finally, compounds in our library target a wide range of human kinases having diverse biological functions (see Supplemental Data Set 2).

Using DSF, we collected temperature denaturation curves for 40 purified kinases in the presence of each one of the 627 compounds in our library (plus vehicle - DMSO; and positive - staurosporine; control). A complete matrix of the thermal shift data is available in Supplemental Data Set 2. A hit was defined as a compound that increased thermal stabilization at least 2x the standard deviation of the DMSO control (Chilton et al., 2017). An example plot of the data and hit identification is depicted in Figure 2A for Os01g01410-cb-001.

**Figure 2.**
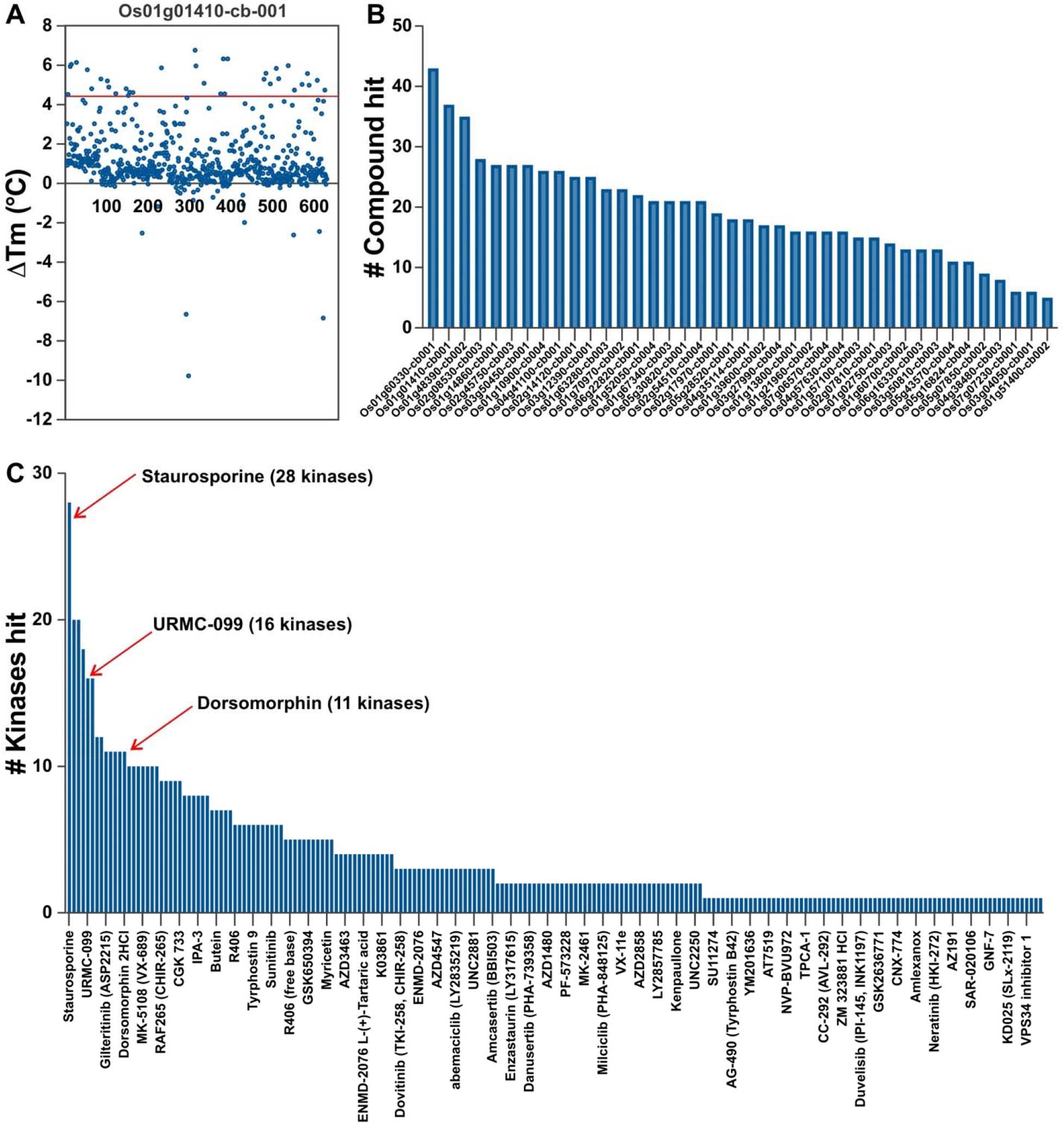
Screening kinases inhibitors against a subset of rice kinases. **(A)** Example thermal shift data set from screening of 627 compounds against Os01g01410-cb-001. Most compounds show no stabilization of the protein, with thermal shift (ΔTm) values near 0 °C. The red line marks 2x the standard deviation of the DMSO control, and hits are defined as compounds that lead to a temperature shift at or above this threshold. **(B)** This bar chart depicts the number of compounds that are classified as hits in the ΔTm assay for each rice kinase screened. Kinases to the left bind many different compounds, while kinases to the right bind only a few of the molecules in the screening set. **(C)** This bar chart provides an indication of the promiscuity of these compounds against this panel of rice kinases. More than a dozen of these compounds (left portion of the bar chart) stabilize 10 or more kinases in the panel, indicating that they are relatively promiscuous, or non-selective. Three of these compounds that are also known to be promiscuous against the human kinome are marked.

As expected, the overall results mirror previous experiments that interrogated a panel of human kinases with a set of kinase inhibitors (Bamborough et al., 2008; Posy et al., 2011; Fedorov et al., 2012; Elkins et al., 2016). In “all versus all” screens, one often identifies promiscuous compounds that bind to many targets, selective compounds that bind very few targets, promiscuous kinase targets that bind a variety of chemotypes, and kinase targets that are more difficult to inhibit and bind relatively few structural classes of inhibitors. Hit rates ranged from a high of 6.8% for Os01g60330-cb001 (43 hits) to a low of 0.8% for Os01g51400-cb002 (5 hits) (Figure 2B).

Figure 2C shows the compounds that qualified as a hit for at least one kinase in the panel, sorted by the number of kinases hit. Three promiscuous human kinase inhibitors (staurosporine, dorsomorphin, URMC-09928) are highlighted that also demonstrate promiscuous binding in this small rice kinase panel. 28 of the 40 rice kinases showed significant stabilization with staurosporine, a very promiscuous human kinase inhibitor. 20 of the compounds stabilized (implying a binding event) 10 or more of these rice kinases screened (Figure 2C). 416 of the compound did not significantly stabilize any of these rice kinases.

Finally, a number of FDA-approved kinase inhibitors are in this screening set, and many show binding to at least one rice kinases (see Supplemental Data Set 3). Some FDA-approved medicines, such as gilteritinib, sunitinib, and vemurafenib, stabilize 5 or more of these rice kinases. A number of quite selective human kinase inhibitors such as the ERBB2 inhibitor lapatinib, EGFR inhibitor gefitinib, and MEK inhibitors trametinib and cobimetinib did not stabilize any of the rice kinases screened.

### Kinase inhibitors affect primary root development in Arabidopsis and rice

From the set of rice kinase inhibitors that were identified by DSF, we selected a group of 37 compounds and tested them for their ability to affect plant development. This subset was chosen to include promiscuous inhibitors targeting several kinases simultaneously and compounds that specifically bind a small group of kinases (Supplemental Table S2). Some of the rice kinases that these compounds bind and likely inhibit, include orthologues of Arabidopsis kinases as BRASSINOSTEROID INSENSITIVE 1 (BRI1) (Li and Chory, 1997; Friedrichsen et al., 2000; Hacham et al., 2011; Kang et al., 2017), SOMATIC EMBRYOGENESIS RECEPTOR-LIKE KINASE 1 and 2 (SERK1 and SERK2) (Du et al., 2012; Gou et al., 2012), CYCLIN-DEPENDENT KINASE F;1 (Takatsuka and Umeda, 2014) and PHYTOSULFOKIN RECEPTOR 1 (PSKR1) (Matsubayashi et al., 2002, 2006) which are known to control root development. Therefore, we decided to first test the activity of these compounds based on their ability to modify the primary root development using Arabidopsis.

Of the 37 compounds tested, 14 caused a significant reduction of primary root length (Figure 3A and 3B and Supplemental Figure S2A and S2B), and only 2 produced a mild increase of elongation compared to the control plants (Figure 3B).

**Figure 3.**
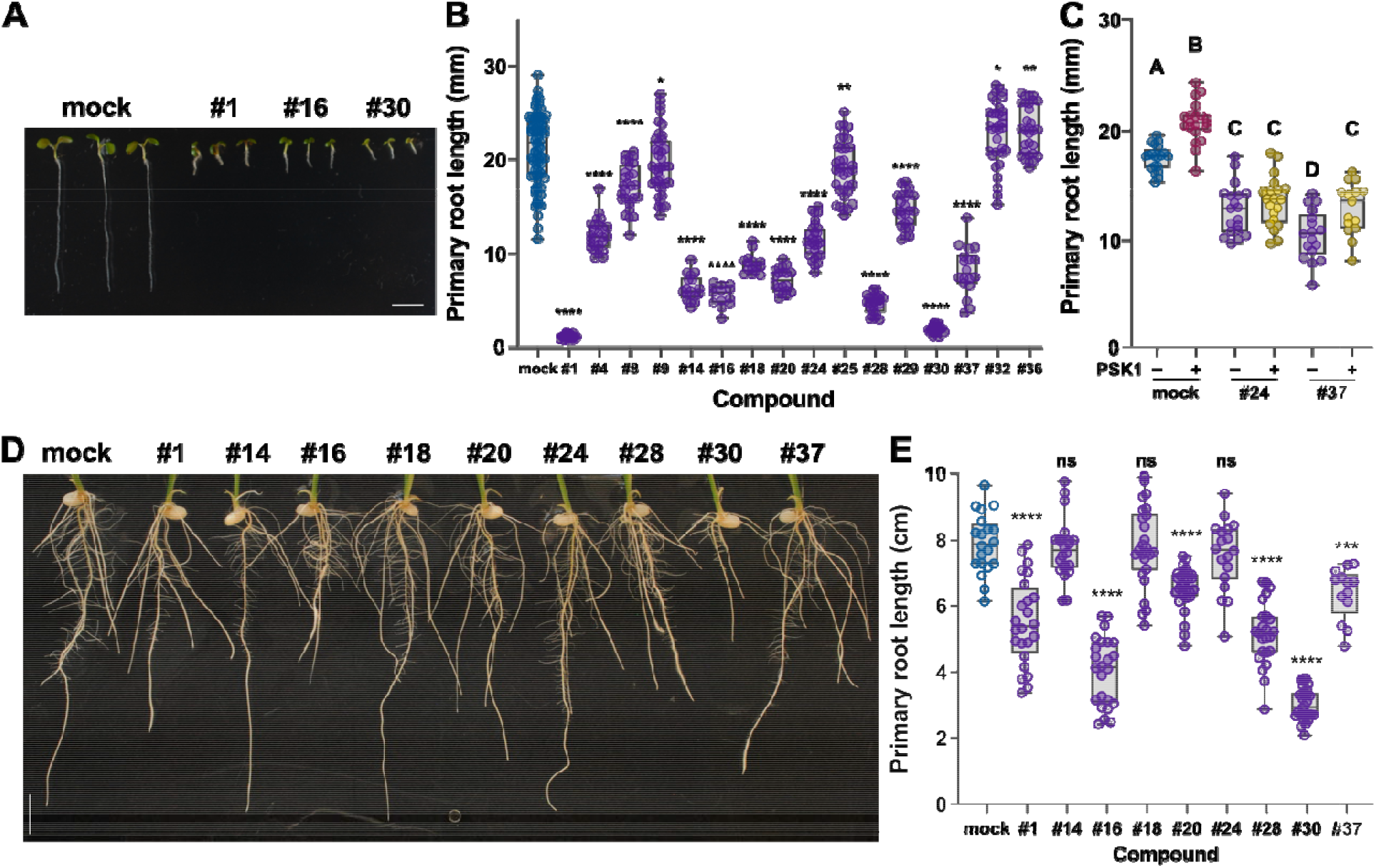
Human kinase inhibitors modify primary root development in Arabidopsis and rice. **(A)** Root growth phenotype 6d after sowing of Col-0 seedlings grown on 1xMS vertical plates with or without 1 µM of the selected kinase inhibitor showing a significant effect on primary root growth. **(B)** and **(C)** Primary root length (mm) 6d after sowing of Col-0 seedlings grown on 1xMS vertical plates with different chemical treatments. In **(B)** plates were prepared with or without 1 µM of the selected kinase inhibitor. In **(C)** we used different combinations of two selected kinase inhibitors that are known to bind the rice orthologue of AtPSKR1 (#24 and #37, 1 µM) and PSK1 (100 nM). **(D)** Root growth phenotype and **(E)** primary root length (cm) 7d after sowing of kitaake seedlings grown on 1xMS vertical plates with or without 1 µM of the selected kinase inhibitor. The data shown in **(B), (C)**, and **(E)** are a box and whisker plot combined with scatter plots, each dot indicates an individual measurement (n=20-30). In **(B)** and **(E)** P values are calculated by two-tailed Student’s t-test (*P ≤ 0.05, **P ≤ 0.01, *** P ≤ 0.001, **** P ≤ 0.0001). In **(C)** different letters indicate significant differences, as determined by ANOVA followed by Tukey’s multiple comparison test (P < 0.05).

We found that compounds #24 (Hesperadin) and #37 (Sitravatinib) each bind the rice kinase (LOC_Os04g57630), which is predicted to be an orthologue of Arabidopsis PSKR1 (AtPSKR1) (see Supplemental Data Set 2), the main receptor of PSK, a small sulfated peptide that positively controls root development (Matsubayashi et al., 2006; Matsuzaki et al., 2010) (Figure 3C). *pskr1* is phenotypically indistinguishable from WT and still responds to synthetic peptide treatment due to the presence of a second PSK receptor (AtPSKR2) that shares almost 50% sequence with AtPSKR1; a double mutant (*pskr1, pskr2*) has shorter roots and is insensitive to PSK treatment (Amano et al., 2007; Pruitt et al., 2017). The short root phenotype observed after chemical treatment may be a consequence of the inhibition of both PSKR1 and PSKR2 (Figure 3B and 3C). In support of this hypothesis, we observed that exogenous addition of the PSK peptide did not rescue the short root growth phenotype in the presence of compound #24. In contrast, a significant response was obtained when combined with compound #37, although the effect was not enough to complement the phenotype to WT (Figure 3C). These results support the hypothesis that these compounds may be totally (compound #24) or partially (compound #37) inhibiting the kinase activity of AtPSKR1 and AtPSKR2. Another possibility is that the lack of response to PSK treatment is a consequence of the inhibition of a different set of kinases, which have a detrimental effect on root growth that is not rescued by PSK treatment..

We next tested the response of rice primary root development to the compounds showing the most significant effects in Arabidopsis. From the 9 compounds tested, 6 caused a reduction in rice primary root development (Figure 3D and 3E).

Three compounds significantly affected Arabidopsis and rice seedling development (#1, #16, and #30) (Figure 3A, 3B, 3D, and 3E). Compound #1, staurosporine, stabilizes 28 kinases in our panel, compound #16 (AD80) stabilizes 10 kinases, and compound #30 (PIK-75) stabilizes 6 kinases. (Supplemental Table S2).

Overall, these results indicate that some of the human kinase inhibitors that can interact with plant kinases based on DSF cause a modification in root growth. Further work is needed to establish structure-activity relationships for individual kinases, verify inhibition of kinase activity in the plant, and build our understanding of the consequences of poly-pharmacology (inhibition of multiple kinases by one compound) on phenotype.

### Multiple sequence alignments suggest that compound RAF265 targets similar regions in human and rice kinases

We next compared the BRAF human kinase with 9 rice kinases stabilized by the same compound RAF265 (CHIR-265). Multiple sequence alignment using the online tool clustal omega (v 1.2.4) revealed that all eleven subdomains indicative of a protein kinase (Hanks et al., 1988) are conserved in the 9 rice kinases and the BRAF human kinase (Supplemental Figure S3). Strikingly, subdomain VI, containing the HRD motif important for catalysis and ending in an invariant Asn involved in substrate binding is particularly well conserved. Furthermore, examination of the BRAF residues involved in binding compound RAF265 according to the co-crystal structure in the PDB (ID 5CT7) (Williams et al., 2015) reveals that these residues are generally highly conserved in the 9 rice kinases (Supplemental Figure S3). These results suggest that RAF265 inhibits the function of both plant and animal kinases in the same manner, as an ATP competitive inhibitor.

## Materials and Methods

### Cloning of rice protein kinase domains into expression vector

Full-length coding DNA (cDNA) clones for the selected rice kinases were used as templates for PCR amplifications. Multiple fragments encompassing the kinase domains (KD) of these genes were amplified and cloned into expression vector pNIC28-Bsa4 (GenBank accession no. EF198106), using ligation-independent cloning (LIC) (Aslanidis and de Jong, 1990; Stols et al., 2002; Gileadi et al., 2008; Burgess-Brown et al., 2014). On average four constructs were designed for each target KD, varying the N- and C-terminal boundaries. T1 phage-resistant *Escherichia coli* Mach-1 cells (Invitrogen, Carlsbard, USA) were used for general cloning. Proteins cloned into pNIC28-Bsa4 vector are fused to an amino-terminal tag of 22 residues (MHHHHHHSSGVDLGTENLYFQ*SM), including a hexahistidine (His6) and a TEV-protease cleavage site. LIC sites are separated by a “stuffer” fragment that contains the *B. subtilis sacB* gene, which allows negative selection on agar plates containing 5% sucrose (Stols et al., 2002). PCR fragments were annealed to the linearized vector through complementary single-stranded regions generated by the T4 DNA polymerase 3’-exonuclease activity. Vector cloning sites were generated by cleavage at two sites by the restriction enzyme *Bsa*I, followed by T4 DNA polymerase treatment in the presence of dGTP. The inserts were treated in the presence of dCTP. Clones were screened by colony PCR and verified by DNA sequencing, using primers specific to the vector: pLIC-F (5’-TGTGAGCGGATAACAATTCC-3’) and pLIC-R (5’-AGCAGCCAACTCAGCTTCC-3’).

### Small-scale test expression

In order to generate expression clones, rice KD constructs were transformed into *E. coli* strains derived from BL21(DE3) and Rosetta 2 (Merck Millipore, Burlington, USA), BL21(DE3)-R3-pRARE2 and BL21(DE3)-R3-lambda-PPase. Strain BL21(DE3)-R3-pRARE2 is a phage-resistant derivative of BL21(DE3) transformed with the pRARE2 plasmid from Rosetta 2 cells, which carries chloramphenicol resistance, while strain BL21(DE3)-R3-lambda-PPase is a phage-resistant derivative of BL21(DE3) transformed with a pACYC-derived plasmid that expresses the bacteriophage-lambda phosphatase as well as three rare tRNAs (Gileadi et al., 2008). Both strains were a kind gift of SGC-Oxford. To find the best constructs and the optimal expression conditions for protein production, all positive clones were evaluated by small-scale test expression followed by IMAC purification from clarified cell lysates. Small-scale test expressions followed the 1-mL expression system described previously (Savitsky et al., 2010; Burgess-Brown et al., 2014). In summary, overnight cultures of expression clones were prepared in 1 mL of Lysogeny broth (LB) medium containing antibiotics (50 µg/mL kanamycin and 34 µg/mL chloramphenicol) in a 96-well deep well block (Sarstedt), and cultures were grown overnight in a microplate shaker (Titramax 101, Heidolph) at 37 °C, shaking at 700 rpm. Overnight cultures (20 μL) were inoculated into 1 mL of Terrific broth (TB) medium containing only kanamycin (50 µg/mL) and incubated at 37 °C, shaking at 900 rpm, until an optical density at 600 nm (OD_600_) of 2-3. Then, expression was induced by adding 0.1 mM IPTG and cultures were incubated overnight at 18 °C, shaking at 700 rpm. Cells were harvested by centrifugation (3,500x g for 20 min) and suspended in 200 µL of lysis buffer [50 mM 4-(2-hydroxyethyl)-1-piperazineethanesulfonic acid (HEPES) pH 7.5, 0.5 M NaCl, 10% glycerol, 10 mM imidazole, 0.5 mM tris-(2-carboxyethyl) phosphine hydrochloride (TCEP)], containing 0.1% dodecyl maltoside (DDM), protease inhibitor cocktail EDTA-free (cat. Number 539134 - Merck Millipore; 1:200), 0.5 mg/mL lysozyme and 50 units/mL benzonase. After freezing the cell suspensions at −80 °C for 20 min, the block was placed in a water bath for approximately 15 min at room temperature, allowing slight thawing. Samples were mixed and an aliquot (3 µL) of the total lysate fraction was removed from each well for future analysis. The lysate was clarified by centrifugation (3,500x g for 10 min) and the supernatant collected in a fresh 96-well deep well block and incubated with 25 µL of pre-equilibrated Ni-sepharose resin (GE Healthcare Life Sciences) in lysis buffer in a microplate shaker (Titramax 101, Heidolph) at 18 °C for 1 hour at 300-400 rpm. The contents of each well were transferred to a 96-well filter plate (Thomson), the resin was washed with 200 µL of wash buffer (50 mM HEPES pH 7.5, 0.5 M NaCl, 10% glycerol, 30 mM imidazole, 0.5 mM TCEP) and centrifuged at 300x g for 1 min. The wash procedure was repeated three more times. Finally, 40 µL of elution buffer (50 mM HEPES pH 7.5, 0.5 M NaCl, 10% glycerol, 300 mM imidazole, 0.5 mM TCEP) was added to each well and proteins were eluted from the resin by centrifugation at 300x g for 3 min. Eluted fractions were analyzed by sodium dodecyl sulfate polyacrylamide gel electrophoresis (SDS-PAGE). The identity of the purified proteins was further confirmed by LS-MS.

### Mid-scale protein expression and purification

Protein expression and purification followed procedures previously described (Tosarini et al., 2018). Briefly, overnight starter cultures were grown in LB medium containing kanamycin (50 µg/mL) and chloramphenicol (34 µg/mL) in an incubator shaker at 37°C, 140 rpm. 5 mL of the starter culture were used to inoculate 500 mL of TB medium supplemented with kanamycin (50 µg/mL). Cells were cultivated at 140 rpm, and 37°C until an OD_600_ ∼1.8. The culture was then transferred to an incubator shaker at 18 °C and kept under 140 rpm. After a 30-min cool-down period, IPTG was added to a final concentration of 0.2 mM. Cells were further cultivated for 16 h at 18°C and 140 rpm. Cells were collected by centrifugation (15 min, 4,000 rpm at 4°C). The pellet was suspended in 2x lysis buffer (1 mL per gram of cells) (1x lysis buffer is 50 mM HEPES pH 7.5, 0.5 M NaCl, 5.0 % (v/v) glycerol, 10 mM imidazole and 1 mM TCEP) supplemented with protease inhibitor cocktail EDTA-free (Merck Millipore; 1:200). Cells were stored at −80°C until use. Cells were lysed by sonication (Sonics Vibra Cell VCX750 ultrasonic cell disrupter) on ice for 5 min (5 sec on, 10 sec off - amplitude = 35%). Polyethyleneimine (PEI - 5% (w/v), pH 7.5) was added to the cell lysate to a final concentration of 0.15%, prior to clarification by centrifugation (45 min, 17,000 rpm, 4°C). Recombinant proteins were enriched from the clarified lysate by gravity-flow metal ion affinity chromatography (IMAC). Chelating Sepharose Fast Flow resin (cat. Number 17057502 - GE Heathcare) was loaded with Ni^2+^ according to the manufacturer’s instructions. A total of 3 ml of Ni^2+^- loaded resin was packed into Econo-Pac columns (cat. Number 7321010 – Bio-Rad) and equilibrated with 3 column volumes (CV) of elution buffer (binding buffer supplemented with 300 mM imidazole - binding buffer is 50 mM HEPES, pH 7.5, 0.5 M KOAc, 10% glycerol, 50 mM arginine and glutamate, 10 mM imidazole, 1 mM TCEP) and 5 CV of binding buffer. Fractions for the flow-through, 10 mM imidazole wash (in binding buffer, 10 CV), 30 mM imidazole wash (in binding buffer, 5 CV) and 300 mM imidazole elution (in binding buffer, 3 CV) were collected and analyzed by 12% SDS-PAGE. Selected IMAC fractions were pooled together and dialyzed (MW cut off 10 kDa) against excess gel filtration buffer (GF buffer is 10 mM HEPES, 0.5 M KOAc, 10% glycerol, 50 mM Arg-Glu, 1 mM TCEP). TEV protease (in a mass ratio of 1:10) was added directly to the dialysis bag. TEV protease treatment was performed overnight at 4 °C. Recombinant proteins lacking the 6His tag were further purified via reverse IMAC using 0.8 ml Ni^2+^- loaded Chelating Sepharose Fast Flow resin packed into poly-prep® chromatography columns (cat. Number 7311550 – Bio-Rad) and prepared as above. Fractions for the flow-through, 10 mM imidazole wash (in GF buffer, 10 CV), 30 mM imidazole wash (in GF buffer, 5 CV) and 300 mM imidazole elution (in GF buffer, 3 CV) were collected and analyzed by 12% SDS-PAGE. Reverse IMAC fractions containing the protein of interest were pooled together and concentrated to a final volume of 5.0 ml. Samples were clarified by centrifugation (10 min at 15,000 rpm and 4 °C) and injected onto a pre-equilibrated Hiload 16/600 Superdex 200 pg (in GF buffer) connected to an AKTApure system (GE Healthcare) set at 0.8 ml/min. Protein samples were concentrated by centrifugation using spin columns (MW cut off of 10 kDa) (cat number UFC501096– Merck Millipore). Protein concentration was estimated by UV using calculated extinction coefficients (42,400 mol/l^-1^.cm^-1^). Protein samples were flash-frozen in a liquid nitrogen bath and stored at −80 °C until use.

### Differential scanning fluorimetry (DSF)

Small molecule screening by DSF was performed as described previously (Niesen et al., 2007; Fedorov et al., 2012). Briefly, the DSF assay was performed in the 96-well format. Purified rice kinase protein was diluted to 2 μM kinase in 100 mM potassium phosphate pH 7.5, 150 mM NaCl, and 10% glycerol supplemented with 5 × SYPRO Orange (Invitrogen). All assay experiments used 19.5 μL of 2 μM kinase and SYPRO Orange mixture. Compounds solubilized in dimethyl sulfoxide (DMSO) were used at a 12.5 μM final concentration, with a 2.5% concentration of DMSO per well. PCR plates were sealed using optically clear films and transferred to a C1000 thermal cycler with CFX-96 RT-PCR head (BioRad). The fluorescence intensity was measured over a temperature gradient from 25 to 95 °C at a constant rate of 0.05 °C/s. Curve fitting and protein melting temperatures were calculated based on a Boltzmann function fitting to experimental data (GraphPad Prism 8). Protein with the addition of 2.5% DMSO was used as a reference. All experiments were carried out in triplicate, and the mean of the ΔTm is reported. Compounds that provided negative values are presented as having a ΔTm of 0 °C.

### Arabidopsis/rice seedling analysis

Seeds from Arabidopsis thaliana accession Col0 and from Oryza sativa ssp. japonica cultivar Kitaake were used in this study. For primary root analysis, Arabidopsis seeds were surface-sterilized in 70% ethanol and then stratified in 0.1% agarose in the dark (4°C) for 2 to 3 days, while rice seeds were dehulled, surface-sterilized in 20% bleach for 30 minutes, and then wash thoroughly with autoclaved water. The seeds were sown on a solid medium containing 1x Murashige and Skoog salt mixture, 1% sucrose (pH 5.8) in 0.3% Gellex (Gellan Gum CAS#71010-52-1 Caisson Laboratories) supplemented with or without 1µM of the selected kinase inhibitor (see Supplemental Table S5 for a description of the compounds tested in this study). The inhibitors were stored as 10µM stocks in DMSO. Plates containing DMSO were used as controls. Synthetic PSK1 is tyrosine-sulfated and was obtained from Pacific Immunology (Ramona, CA, USA). The peptide was stored as 1 µM stocks in water ddH_2_O. The top half of the Petri dish was sealed with Micropore tape to allow gas exchange and plates were placed vertically for 6 days in chambers with 16-h-light/8-h-dark photoperiod at 21°C for Arabidopsis and for 7 days in incubators with 14-h-light/10-h-dark photoperiod at 28°C/24 °C for rice. The seeds germinated properly in the plates from all the inhibitors, discarding any effect these compounds might have on seed germination Plates were photographed, and the root length was measured with Fiji (Schindelin et al., 2012).

## Supporting information

Supplemental Data Set 3

Supplemental Data Set 2

Supplemental Data Set 1

## Acknowledgments

This work was supported by the Foundation for Food and Agricultural Research (FFAR) grant #534683 to PCR.

PZR and RMC acknowledge support from CAPES (Coordenação de Aperfeiçoamento de Pessoal de Nível Superior; grant number: 88887.136386/2017-00), FAPESP (Fundação de Amparo à Pesquisa do Estado de São Paulo; grant numbers: 2013/50724-5 and 2014/50897-0) and CNPq (Conselho Nacional de Desenvolvimento Científico e Tecnológico; grant number: 465651/2014-3). PZR received a CAPES post-doctoral fellowship (88887.136432/2017-00). MFE is a Latin American Fellow in the Biomedical Sciences, supported by the Pew Charitable Trusts.

Christian Rogers at the Sainsbury Laboratory for gene synthesis.

## Author Contributions

MF Ercoli: Root assays, data analysis.

PZ Ramos: Cloning into expression vector, small-scale test expression, data analysis

Rashmi Jain: Selection of kinases and Figure 1 (Phylogenetic tree), data analysis

J Pilotte: Protein production, DSF screening

Oliver Xiaoou Dong: Coordinated the *de novo* synthesis of the 129 rice kinase genes.

Ty Thompson: Root assays

C Wells: DSF screening, data analysis

JM Elkins: Kinase domain analysis and expression construct design.

AM Edwards: project conceptualization.

RM Couñago: Manuscript writing, data analysis

PC Ronald: project conceptualization, manuscript writing

DH Drewry: project conceptualization, manuscript writing, data analysis

All: manuscript editing

