## Supplemental Figures for "An open source plant kinase chemogenomics set"

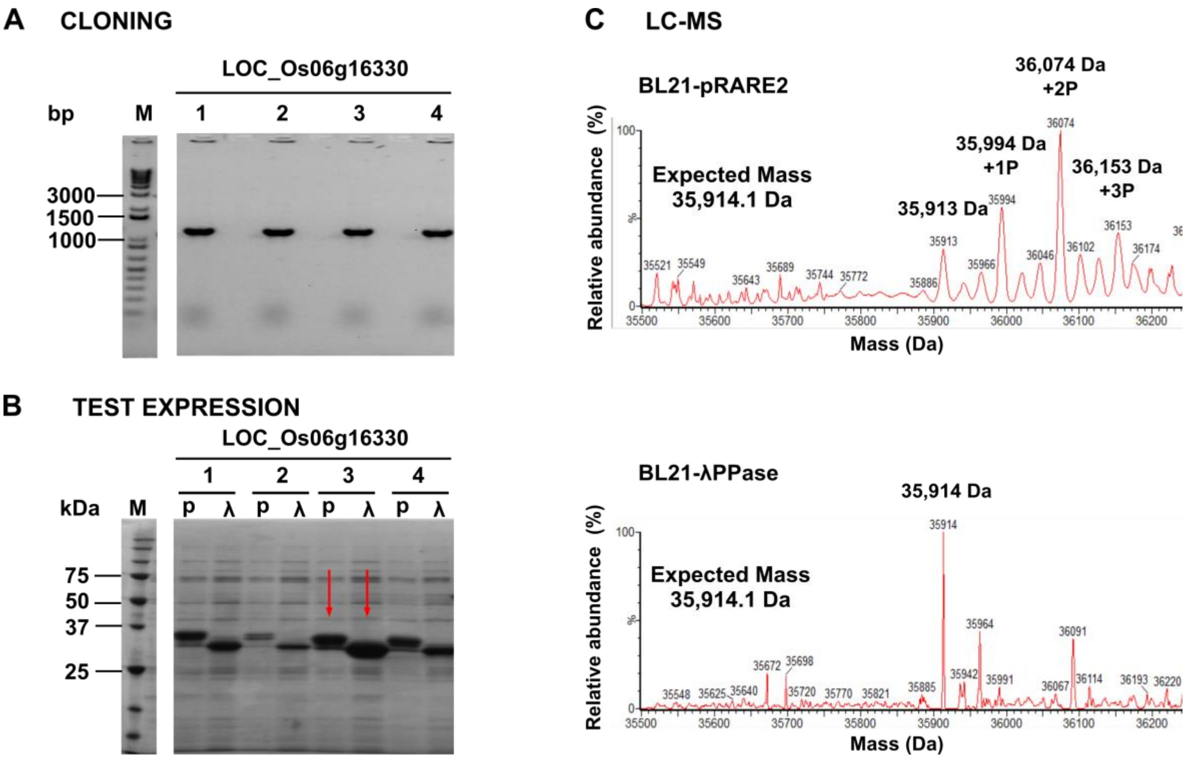


**Supplemental Figure S1. Cloning and small-scale test expression of LOC_Os06g16330 kinase domain (KD).** (**A**) Agarose gel showing amplicons from positive clones amplified from bacterial colonies by PCR. Four different constructs were designed for this rice kinase. M: molecular weight marker (1 Kb Plus DNA Ladder, Invitrogen). (**B)** SDS-PAGE analysis of eluted fractions obtained from small-scale test expression in both BL21(DE3)-R3-pRARE2 (p) and BL21(DE3)-R3-lambda-PPase (λ) strains. M: molecular weight marker (Precision Plus Protein Unstained Protein Standards, Bio-Rad). (**C)** Liquid chromatography-mass spectrometry (LC-MS) analysis for LOC_Os06g16330 KD purified from small-sale test expression (construct 3 – indicated in panel B). Deconvoluted mass/charge spectra are shown for proteins expressed in BL21(DE3)-R3-pRARE2 (top) and BL21(DE3)-R3-lambda-PPase (bottom) strains. Expected and observed mass values are indicated. In addition to the correct mass, three phosphorylation states were noticed when the protein was not co-expressed with Lambda Protein Phosphatase (top).

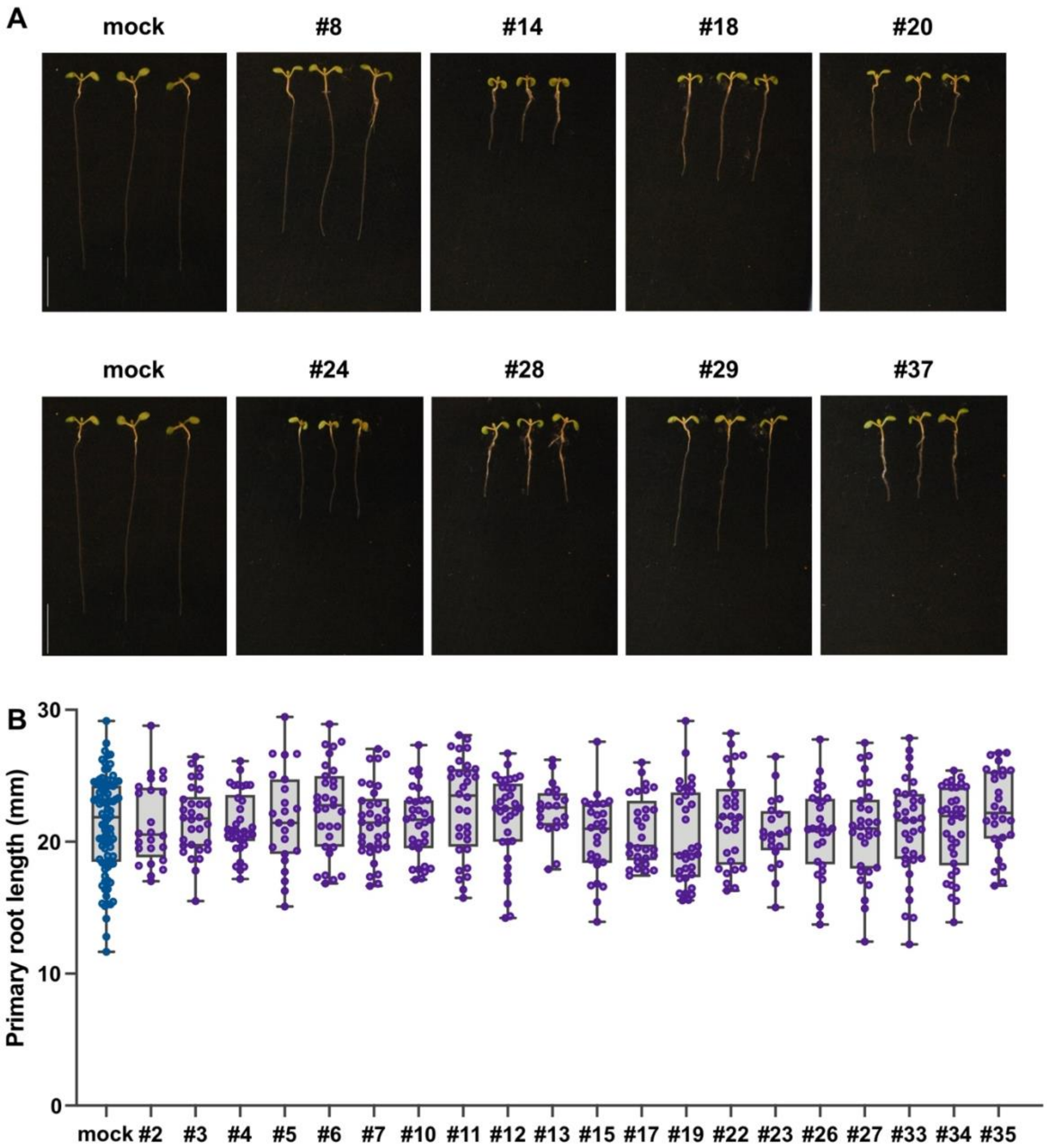


**Supplementary Figure S2. Root growth phenotypes in Arabidopsis plants treated with human kinase inhibitors.**
**(A)** Root growth phenotype 6d after sowing of Col-0 seedlings grown on 1xMS vertical plates with or without 1 µM of the selected kinase inhibitor showing a significant effect on primary root growth. **(B)** Primary root length (mm) 6d after sowing of Col-0 seedlings grown on 1xMS vertical plates with or without 1 µM of the kinase inhibitors without significant effects on Arabidopsis root growth. The data shown in **(B)** are a box and whisker plot combined with scatter plots, each dot indicates an individual measurement (n=20-30).

CLUSTAL O(1.2.4) multiple sequence alignment

```
                                        I     * *           II                           III*
LOC_Os05g43570.2:116-420         ELLTIIGRGAFGEVRLCREKASKNVYAMKKLKKSEML--------RRGQVEHVKAERNLL     52
LOC_Os06g22820.1:24-419          EVLGRAGSGAYADVYRGRRRSDGAPVALKEVHDAVS----------------ARREADAL     44
P15056-BRAF_HUMAN:457-717        TVGQRIGSGSFGTVYKGKW-HG--DVAVKMLNVTAP---------TPQQLQAFKNEVGVL     48
LOC_Os02g02780.1:305-557         KFGTKVASGSNGDLFRGSY-CS-QDVAIKVVRPERI---------SADMYRDFAQEVYIM     49
LOC_Os01g48390.2:352-620         DFSNIVASYPHYTVYKGTL-SSGVEIAVVSTVIAT------NKDWSKHSEGRFRKKIDLL     53
LOC_Os01g60330.1:343-620         ASAEVLGKGSYGTTYKAVL-EDGTTVVVKRLKEVV------------VGKKDFEQQMEIV     47
LOC_Os01g63280.2:138-414         SEQNRIGLGNFGTVYKGKL-RDGSIIAVKRATKNMYD---------RHLSEEFRSEIQTL     50
LOC_Os01g02750.1:106-395         RFKVKVGQGGFGSVYRGEL-PNGVPVAVKMLENPK------------GEGDEFINEVATI     47
LOC_Os01g13800.1:690-981         DEDNVIGSGASGKVYKAVL-SNGEVVAVKKLWGLKKGTDVENGGEGSTADNSFEAEVKTL     59
LOC_Os01g21960.1:193-457         SKDNILGEGGYGVVYRGQL-INGTPVAVKKLLNNL-----------GQAEKEFRVEVEAI     48
                                       .              .    .:                           :   :

                                         ** IV            * * *
LOC_Os05g43570.2:116-420         AEV-DSAFIVKLYYSFQ--DEEYLYLIMEYLPGGDMMTLLMRKDT------LTEDEARFY     103
LOC_Os06g22820.1:24-419          LAAAPSRHVVALLDHFPGGDHDDDVLVLEWLPLDLSAVVRAAAAA--RPSALPAAQRKRW     102
P15056-BRAF_HUMAN:457-717        RKT-RHVNILLFMGYST-KP--QLAIVTQWCEGSSLYHHLHIIE-----TKFEMIKLIDI     99
LOC_Os02g02780.1:305-557         RKV-RHRNVVQFIGACTRQP--NLYIVTDFMSGGSLHDYLHKKN-----NSFKLSEILRV     101
LOC_Os01g48390.2:352-620         SRI-NHKNFINLLGYCEEENPFMRMMVLEYAPNGTLYEHLHVE----GFDHIDWNGRMRV     108
LOC_Os01g60330.1:343-620         GRVGQHQNVVPLRAYYYSKD--EKLLVYDYIPSGSLAVVLHGNKA-TGKAPLDWETRVKI     104
LOC_Os01g63280.2:138-414         SKV-EHLNLVKFLGYLEHED--ERLILVEYVNNGSLREHLDGLR----GEPLEFSQRLNI     103
LOC_Os01g02750.1:106-395         GRI-HHANIVRLLGFCSEGT--RRALIYEYIPNDSLEKYIFSHDSNTSQELLVPSKMLDI     104
LOC_Os01g13800.1:690-981         GKI-RHKNIVKLWCSCTHND--TKLLVYEYMPNGSLGDVLHS--SK--AGLLDWSTRYKI     112
LOC_Os01g21960.1:193-457         GHV-RHKNLVRLLGYCVEGT--QRMLVYEYVNNGNLEQWLHGAMSH--RGSLTWEARVKI     103
                                         .: :             :: ::   .                 :

                                        V  *           VI     *        *VII
LOC_Os05g43570.2:116-420         IAETVLAIESIHK---HSYIHRDIKPDNLLLDRSGHLKLSDFGLCKPLDSSNFPNLNEPD     160
LOC_Os06g22820.1:24-419          MLQVLEGVAACHS---AGVVHRDLKPANLLISEDGVLKVADLGQARILQETGTYQG-MHP     158
P15056-BRAF_HUMAN:457-717        ARQTAQGMDYLHA---KSIIHRDLKSNNIFLHEDLTVKIGDFGLATVKSR-WS--G-SHQ     152
LOC_Os02g02780.1:305-557         ATDISKGMNYLHQ---NNIIHRDLKTANLLMDENKVVKVADFGVARVKDQ-S------GV     151
LOC_Os01g48390.2:352-620         IMGVAYCIQHMHE-LNPSITHPDLHSSAILLSEDGAAKVADMSVWQEVISKGK--M-PKN     164
LOC_Os01g60330.1:343-620         SLGVARGIAHLHAEGGGKFIHGNLKSSNILLSQNLDGCVSEFGLAQLMTIPP--------     156
LOC_Os01g63280.2:138-414         AIDIVHAVSYLHGYTDHPIIHRDIKSSNILLTDQLRAKVADFGFARLAPDNTE--A-THV     160
LOC_Os01g02750.1:106-395         ALGIARGMEYLHQGCNQRILHFDIKPNNILLDYNFSPKISDFGLAKLCARDQS----IVT     160
LOC_Os01g13800.1:690-981         ALDAAEGLSYLHHDYVPAIVHRDVKSNNILLDAEFGARVADFGVAKVVEATVR--G-PKS     169
LOC_Os01g21960.1:193-457         LLGTAKALAYLHEAIEPKVVHRDIKSSNILIDDDFDAKVSDFGLAKLLGAGKS-----HV     158
                                        :   *        * :::   :::  .    :.::.

                                          VIII
LOC_Os05g43570.2:116-420         YTSTKGTKPLPDSSSRLSSSALKRTQ------------QEQL------------------     190
LOC_Os06g22820.1:24-419          YEQSSGVEPWVSQQRAVL-HGVKENHPSHDSETQTGQEPERLTAADYLHEMDQLRAKSTH     217
P15056-BRAF_HUMAN:457-717        FEQLSGSILWMAPEVIRM------------------------------------------     170
LOC_Os02g02780.1:305-557         MTAETGTYRWMAPEVIE-------------------------------------------     168
LOC_Os01g48390.2:352-620         DDI-----------VDHH------------------------------------------     171
LOC_Os01g60330.1:343-620         --APARLVGYRAPEVL--------------------------------------------     170
LOC_Os01g63280.2:138-414         STMVKGTAGYVDPEYM--------------------------------------------     176
LOC_Os01g02750.1:106-395         LTAARGTMGYIAPELYSR------------------------------------------     178
LOC_Os01g13800.1:690-981         MSVIAGSCGYIAPEYA--------------------------------------------     185
LOC_Os01g21960.1:193-457         TTRVMGTFGYVAPEYA--------------------------------------------     174

LOC_Os05g43570.2:116-420         --------------------------------SHWQ------KNRRMLAYSTVGTPDYIA     212
LOC_Os06g22820.1:24-419          GDVDKMSLQDGNASCLATCSTADIDDDPFRASYSYDAEEGMLEEESGAFTSCVGTRWFRA     277
P15056-BRAF_HUMAN:457-717        ------------------------------------------------------------     170
LOC_Os02g02780.1:305-557         ------------------------------------------------------------     168
LOC_Os01g48390.2:352-620         ------------------------------------------------------------     171
LOC_Os01g60330.1:343-620         ------------------------------------------------------------     170
LOC_Os01g63280.2:138-414         ------------------------------------------------------------     176
LOC_Os01g02750.1:106-395         ------------------------------------------------------------     178
LOC_Os01g13800.1:690-981         ------------------------------------------------------------     185
LOC_Os01g21960.1:193-457         ------------------------------------------------------------     174
```

```
                                              IX        X
LOC_Os05g43570.2:116-420         PEVLL-KKGYGMECDWWSLGAIMYEMLVGYPPFYSEDPMSTCRKIVNWRSHLK---FPEE     268
LOC_Os06g22820.1:24-419          PELLYGSTNYGQEVDLWSLGCILAELFNLEPIFPGTSDIDQIGRIISVLGNITEETFPGC     337
P15056-BRAF_HUMAN:457-717        ----QDKNPYSFQSDVYAFGIVLYELMTGQLPYSN-INNRDQ-----IIF-MVGR----G     215
LOC_Os02g02780.1:305-557         ------HKPYDHKADVFSFGIVLWELLTGKIPYEY-LTPLQA-----AIG-VVQK----G     211
LOC_Os01g48390.2:352-620         ----E-PVSADPAGNVCSFGLLMLEIISGRPPYSE-HKGS---LANLAMECIKDDRNISC     222
LOC_Os01g60330.1:343-620         ----E-TKKPTQKSDVYSFGVLVLEMLTGKAPLRSPGREDSIEHLPRWVQSVVREEWTAE     225
LOC_Os01g63280.2:138-414         ----R-TNQLTDRSDVYSFGVLLVELLTGRRPIE-RGRGRHQRLTTQWALRKCRDGDAVV     230
LOC_Os01g02750.1:106-395         ----N-FGEISYKSDVYSFGMLVLEMVSGRRNSD-PSVESQN--EVYFPECIYEQVTTGR     230
LOC_Os01g13800.1:690-981         ----Y-TLRVNEKSDIYSFGVVLLELVTGKPPVD-PEF--GEKDLVKWVCSTIDQKGVEH     237
LOC_Os01g21960.1:193-457         ----N-TGLLNEKSDIYSFGVVLLEAITGRDPVD-YGRPANEVNLVDWLKMMVASRRSEE     228
                                              :  ::* :: * .

                                                                              XI
LOC_Os05g43570.2:116-420         ------AKLSP------------------EAKDLISKLLCNVEQ---RLGTKGAHE----     298
LOC_Os06g22820.1:24-419          SNLPDYNKIFFNKVEKPIGLEACLPDRSASEVSIIKRLLCY--DPTKRASAADL------     389
P15056-BRAF_HUMAN:457-717        YLSPDLSKV--RS--------NCP------KAMKRLMAECLKKKRDERPLFPQILASIEL     259
LOC_Os02g02780.1:305-557         LR-PTI-----PK--------DTH------PKLSELLQKCWHRDPAERPDFSQILEILQR     251
LOC_Os01g48390.2:352-620         LLDPTLKTHK--E--------NEL------EIICELIQECIQSDPKKRPGMREVTTRLRE     266
LOC_Os01g60330.1:343-620         VFDVDLLRHPNIE--------DEM------VQMLQVAMACVAAPPDQRPKMDEVIRRIVE     271
LOC_Os01g63280.2:138-414         AMDARMRRTSAVV--------AAM------EKVMALAAECTAPDRAARPAMRRCAEVLWS     276
LOC_Os01g02750.1:106-395         DLELGREMTQEEK--------ETM------RQLAIVALWCIQWNPKNRPSMTKVVNMLTG     276
LOC_Os01g13800.1:690-981         VLDSKLDMT-F-K--------DEI------NRVLNIALLCSSSLPINRPAMRRVVKMLQE     281
LOC_Os01g21960.1:193-457         VVDPTIETR-PST--------RAL------KRALLTALRCVDPDSEKRPKM---------     264
                                                                        *       *

LOC_Os05g43570.2:116-420         -------KAHPWF 304
LOC_Os06g22820.1:24-419          -------LNDPYF 395
P15056-BRAF_HUMAN:457-717        L------------ 260
LOC_Os02g02780.1:305-557         L------------ 252
LOC_Os01g48390.2:352-620         VL----------- 268
LOC_Os01g60330.1:343-620         IRNSYS------- 277
LOC_Os01g63280.2:138-414         ------------- 276
LOC_Os01g02750.1:106-395         RLQNLHVPPKPFF 289
LOC_Os01g13800.1:690-981         VRAEATRPRL--- 291
LOC_Os01g21960.1:193-457         ------------- 264
```

**Supplementary Figure 3. Multiple sequence alignment showing sequence similarity between human and rice kinases hit by the same compounds.** All 11 subdomains indicative of a protein kinase are highlighted with roman numerals above the alignment. BRAF Residues involved in binding compound RAF265 are marked with red stars above the alignment, excluding the invariant DFG motif and Lys-Glu bridge which are involved in binding RAF265 but are highly conserved in all protein kinases, indicating that BRAF residues involved in binding RAF265 show a high degree of conservation across the 9 rice kinases that bind RAF265, suggesting that RAF265 is also an ATP-competitive inhibitor of the rice kinases.

| | | | | |
|---|---|---|---|---|
| AAC49123 (Xa21) | LOC_Os02g53720 | LOC_Os06g16070 | LOC_Os06g16070 | LOC_Os12g01200 |
| LOC_Os01g01410 | LOC_Os02g54510 | LOC_Os06g16330 | LOC_Os06g16330 | LOC_Os12g01510 |
| LOC_Os01g02730 | LOC_Os02g57080 | LOC_Os06g22820 | LOC_Os06g22820 | LOC_Os12g07990 |
| LOC_Os01g02750 | LOC_Os02g57440 | LOC_Os06g41960 | LOC_Os06g41960 | LOC_Os12g13300 |
| LOC_Os01g10900 | LOC_Os03g02680 | LOC_Os06g45240 | LOC_Os06g45240 | LOC_Os12g37980 |
| LOC_Os01g13800 | LOC_Os03g04050 | LOC_Os07g04220 | LOC_Os07g04220 | LOC_Os12g40279 |
| LOC_Os01g14860 | LOC_Os03g12390 | LOC_Os07g04820 | LOC_Os07g04820 | LOC_Os12g41090 |
| LOC_Os01g16230 | LOC_Os03g14710 | LOC_Os07g06570 | LOC_Os07g06570 | LOC_Os12g41260 |
| LOC_Os01g20880 | LOC_Os03g14840 | LOC_Os07g07230 | LOC_Os07g07230 | LOC_Os12g43640 |
| LOC_Os01g21960 | LOC_Os03g18170 | LOC_Os07g37810 | LOC_Os07g37810 | |
| LOC_Os01g36080 | LOC_Os03g27990 | LOC_Os07g38530 | LOC_Os07g38530 | |
| LOC_Os01g38840 | LOC_Os03g31044 | LOC_Os08g05290 | LOC_Os08g05290 | |
| LOC_Os01g39600 | LOC_Os03g47470 | LOC_Os08g10320 | LOC_Os08g10320 | |
| LOC_Os01g46030 | LOC_Os03g50230 | LOC_Os08g17410 | LOC_Os08g17410 | |
| LOC_Os01g48390 | LOC_Os03g50450 | LOC_Os08g34380 | LOC_Os08g34380 | |
| LOC_Os01g49920 | LOC_Os03g50810 | LOC_Os08g40280 | LOC_Os08g40280 | |
| LOC_Os01g51400 | LOC_Os03g51020 | LOC_Os08g40990 | LOC_Os08g40990 | |
| LOC_Os01g52050 | LOC_Os03g57940 | LOC_Os09g16980 | LOC_Os09g16980 | |
| LOC_Os01g56580 | LOC_Os03g58750 | LOC_Os09g17630 | LOC_Os09g17630 | |
| LOC_Os01g57100 | LOC_Os03g60150 | LOC_Os09g23570 | LOC_Os09g23570 | |
| LOC_Os01g59560 | LOC_Os04g20680 | LOC_Os09g31210 | LOC_Os09g31210 | |
| LOC_Os01g60330 | LOC_Os04g24510 | LOC_Os09g33630 | LOC_Os09g33630 | |
| LOC_Os01g60700 | LOC_Os04g29770 | LOC_Os09g39860 | LOC_Os09g39860 | |
| LOC_Os01g63280 | LOC_Os04g35114 | LOC_Os10g05160 | LOC_Os10g05160 | |
| LOC_Os01g66020 | LOC_Os04g38480 | LOC_Os10g10540 | LOC_Os10g10540 | |
| LOC_Os01g67340 | LOC_Os04g41100 | LOC_Os10g38800 | LOC_Os10g38800 | |
| LOC_Os01g70970 | LOC_Os04g46320 | LOC_Os10g38960 | LOC_Os10g38960 | |
| LOC_Os02g01730 | LOC_Os04g57630 | LOC_Os11g01200 | LOC_Os11g01200 | |
| LOC_Os02g02140 | LOC_Os05g07850 | LOC_Os11g02305 | LOC_Os11g02305 | |
| LOC_Os02g02780 | LOC_Os05g11140 | LOC_Os11g06140 | LOC_Os11g06140 | |
| LOC_Os02g05730 | LOC_Os05g15720 | LOC_Os11g10310 | LOC_Os11g10310 | |
| LOC_Os02g07810 | LOC_Os05g16824 | LOC_Os11g11490 | LOC_Os11g11490 | |
| LOC_Os02g08530 | LOC_Os05g28520 | LOC_Os11g17080 | LOC_Os11g17080 | |
| LOC_Os02g14120 | LOC_Os05g30820 | LOC_Os11g25860 | LOC_Os11g25860 | |
| LOC_Os02g17970 | LOC_Os05g34390 | LOC_Os11g29510 | LOC_Os11g29510 | |
| LOC_Os02g21700 | LOC_Os05g40050 | LOC_Os11g35220 | LOC_Os11g35220 | |
| LOC_Os02g39560 | LOC_Os05g43570 | LOC_Os11g39370 | LOC_Os11g39370 | |
| LOC_Os02g41580 | LOC_Os05g50120 | LOC_Os11g40550 | LOC_Os11g40550 | |
| LOC_Os02g42160 | LOC_Os05g51400 | LOC_Os11g42440 | LOC_Os11g42440 | |
| LOC_Os02g45750 | LOC_Os06g05830 | LOC_Os11g44660 | LOC_Os11g44660 | |

**Supplemental Table S1. List of the 129 rice protein kinases selected for this study. Names are locus identifiers according to the Rice Genome Annotation Project (http://rice.uga.edu/), except for AAC49123 (Xa21), which is the GenBank ID.**

| Compound number | Compound name | # of kinases hit |
|---|---|---|
| 1 | staurosporine | 28 |
| 2 | NCB-0846 | 18 |
| 3 | SGI-7079 | 20 |
| 4 | SB216763 | 12 |
| 5 | LY2874455 | 20 |
| 6 | Sennoside B | 11 |
| 7 | Shikonin | 11 |
| 8 | RAF265 (CHIR-265) | 9 |
| 9 | AS-252424 | 9 |
| 10 | URMC-099 | 16 |
| 11 | THZ1 | 10 |
| 12 | UNC2025 | 16 |
| 13 | AST-1306 | 9 |
| 14 | MK-5108 (VX-689) | 10 |
| 15 | SU6656 | 8 |
| 16 | AD80 | 10 |
| 17 | AT9283 | 10 |
| 18 | vemurafenib (PLX4032, RG7204) | 6 |
| 19 | GW5074 | 8 |
| 20 | GSK650394 | 5 |
| 21 | NT157 | 10 |
| 22 | dorsomorphin | 11 |
| 23 | BX-912 | 10 |
| 24 | hesperadin | 12 |
| 25 | BX-795 | 8 |
| 26 | Tyrphostin 9 | 6 |
| 27 | dovitinib (TKI-258) | 11 |
| 28 | GZD824 | 6 |
| 29 | BIO | 5 |
| 30 | PIK-75 | 5 |
| 31 | BI-78D3 | 5 |
| 32 | 7,8-dihydroxyflavone | 7 |
| 33 | (+)-epigallocatechin gallate | 5 |
| 34 | butein | 7 |
| 35 | myricetin | 5 |
| 36 | gilteritinib (ASP2215) | 11 |
| 37 | sitravatinib (MGCD516 | 6 |

**Supplemental Table S2. List of rice kinase inhibitors selected to perform root growth assay and the rice kinases they bind based on DSF results**
